# Targeted Drug Repurposing Against the SARS-CoV-2 E channel Identifies Blockers With *in vitro* Antiviral Activity

**DOI:** 10.1101/2021.02.24.432490

**Authors:** Prabhat Pratap Singh Tomar, Miriam Krugliak, Isaiah Tuvia Arkin

## Abstract

It is difficult to overstate the impact that COVID-19 had on humankind. The pandemic’s etiological agent, SARS-CoV-2, is a member of the Coronaviridae, and as such, is an enveloped virus with ion channels in its membrane. Therefore, in an attempt to provide an option to curb the viral spread, we searched for blockers of its E protein viroporin. Using three bacteria-based assays, we identified eight compounds that exhibited activity after screening a library of ca. 3000 approved-for-human-use drugs. Reassuringly, analysis of viral replication in tissue culture indicated that most of the compounds could reduce infectivity to varying extents. In conclusion, targeting a particular channel in the virus for drug repurposing may increase our arsenal of treatment options to combat COVID-19 virulence.

**Significance Statement:** The goal of our study was to expand the treatment arsenal against COVID-19. To that end, we have decided to focus on drug therapy, and as a target - the E protein, an ion channel in the virus. Ion channels as a family are excellent drug targets, but viral channels have been underexploited for pharmaceutical point intervention. To hasten future regulatory requirements and focus the chemical search space, we screened a library of ca. 3000 approved-for-human-use drugs using three independent bacteria-based assays. Our results yielded eight compounds, which were subsequently tested for antiviral activity in tissue culture. Gratifyingly, most compounds were able to reduce viral replication, and as such, both validate our approach and potentially augment our anti-COVID tool kit.

---

At the end of 2019, a new respiratory disease engulfed much of the globe. Approximately 100 million people were identified as carrying the virus in a year, with a mortality rate exceeding 2% (1). The pandemic’s etiological agent was quickly identified as a new coronavirus (2,3) and was found to be very similar to the virus that caused the SARS epidemic in 2002/3 (4, 5). Accordingly, the virus was named: SARS-CoV-2 (6).

As a member of the Coronaviridae, SARS-CoV-2 is an enveloped virus and contains several proteins in its membrane. One of the membrane constituents is the E protein, which has been implicated in viral assembly, release, and pathogenesis, based on studies in other coronaviruses (7). Notably, E proteins were shown to be essential for viral infectivity (8), and attenuated viruses that lack them were suggested to serve as vaccine candidates (9–12).

Functionally, E proteins from several coronaviruses, including the very similar SARS-CoV-1, were shown to possess cation-selective channel activity (13–15). Consequently, we have recently confirmed that the E protein from SARS-CoV-2 is also a channel using bacteria-based assays (16).

As a protein family, ion channels serve as excellent targets for pharmaceutical point intervention. For example, chemicals that manipulate ion channels are used to treat many diseases such as cystic fibrosis, epilepsy, arrhythmia, neurodegenerative diseases, hypertension, angina, and more (17).

Ion channels in viruses have also been suggested to serve as attractive drug targets (18). A prominent example is the antiflu aminoadamantane drugs (19) that target influenza’s M2 protein (20) by blocking its channel activity (21). Regrettably, wide-spread viral resistance has rendered aminoadamantanes ineffective (22).

Considering the above, we have decided to search for blockers against the SARS-CoV-2 E protein channel as a potential approach to curb infectivity. In preliminary studies employing a small library of channel blockers, we identified two low-affinity inhibitors of the protein (16), motivating further screening efforts. Therefore, in the current study, we focussed our search on a significantly larger library of ca. 3000 approved-for-use compounds. Drug repurposing as such minimizes the chemical space to explore and may potentially hasten any future regulatory steps.

## Results

The approach that we have taken involves two components employed in serial fashion. We started by screening a repurposed drug library using three bacteria-based assays. Subsequently, the antiviral activity of the hits was tested in tissue culture studies.

### Library screening

Our screening strategy employed the analysis of bacteria that heterologously express the E protein. Subsequently, the effect of different chemicals on the channel may be examined by monitoring their impact on the bacteria.

To ensure proper membrane reconstitution, we used the MBP Fusion and Purification System (New England BioLabs, Ipswich, MA) in which the E protein was fused to the carboxy terminus of the maltose-binding protein. This system has been used to express and study numerous other viral ion channels successfully (23–26). Finally, we have recently shown that SARS-CoV-2 E protein is also expressed in functional form utilizing this construct (16).

#### Negative assay

The first assay that we used is one in which the E protein is expressed at elevated levels in the bacteria. Consequently, the bacteria experience severe growth retardation due to excessive membrane permeabilization caused by the viral channel. In other words, the viral channel impacts the bacteria **negatively**. As a result, blockers of the viral channel may be identified due to their ability to revive bacterial growth.

Using the above approach, we screened ca. 3000 chemicals from the drug repurposing library of MedChem Express (Monmouth Junction, NJ). Specifically, bacterial cultures were grown overnight in 96-well plates, and the impact of each chemical in the library at a concentration of 50 *µ*M was tested individually. Finally, any hit was then analyzed at several different concentrations to obtain a dose-response curve.

The results shown in Figure 1a indicate that the following eight chemicals are able to revive bacterial growth to varying extents: 5-Azacytidine (+84%), Plerixafor (+173%), Mebrofenin (+263%), Mavorixafor (trihydrochloride) (+302%), Plerixafor (octahydrochloride) (+137%), Cyclen (+359%), Kasugamycin (hydrochloride hydrate) (+141%), and Saroglitazar Magnesium (+120%). The values in parenthesis are the growth enhancement of each chemical at 50 *µ*M relative to untreated bacteria. Finally, see Supplementary Figures 1-8 for detailed chemical structures and raw growth curves of each of the compounds.

**Fig. 1.**
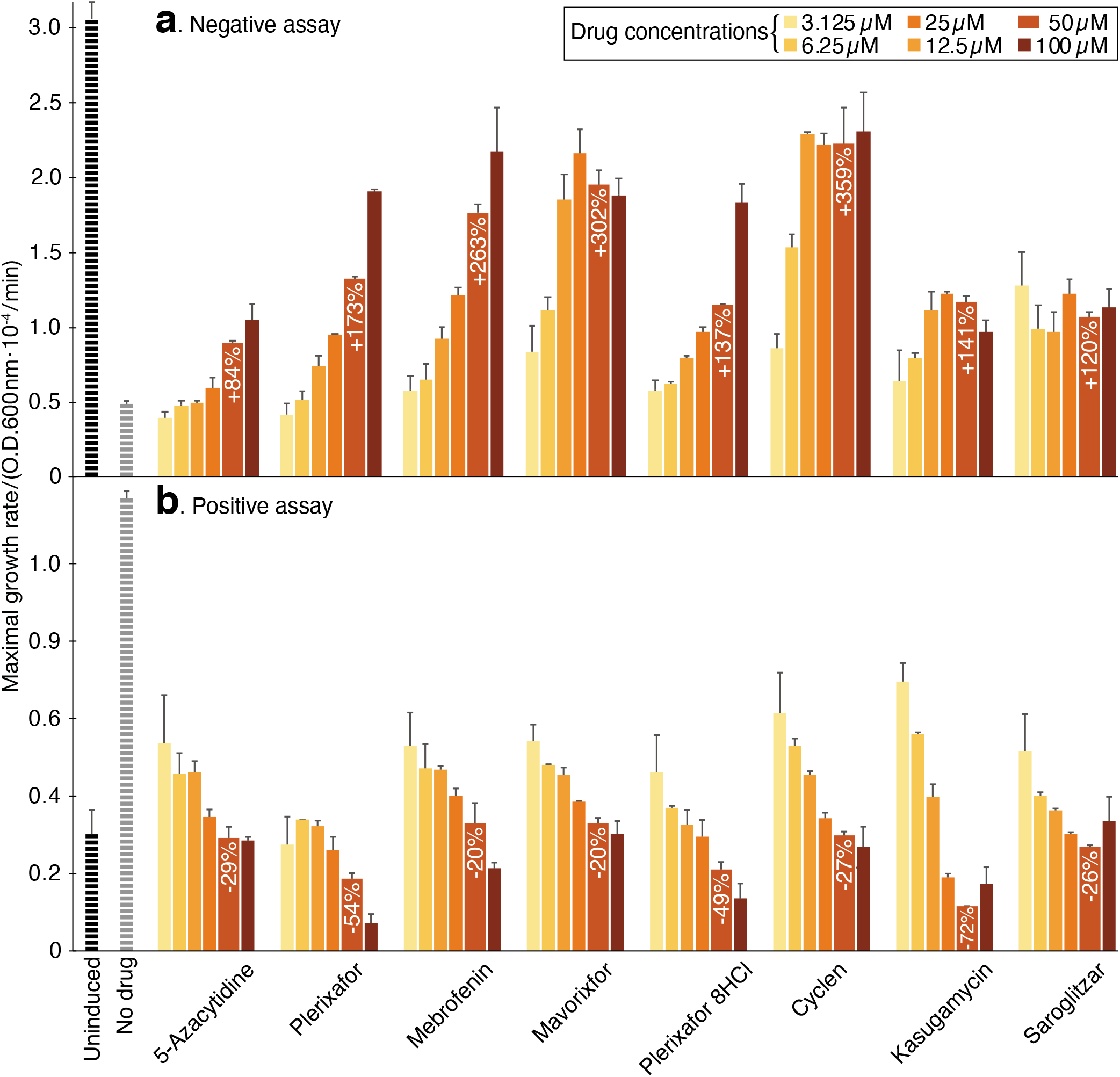
Compound screening results using the negative and positive assays. a. Negative assay in which SARS-CoV-2 E protein is expressed at an elevated level (induction with 100 *µ*M [*β*-D-1-thiogalactopyranoside]) and is therefore deleterious to bacteria. In this instance, inhibitory drugs enhance bacterial growth. b. Positive assay in which SARS-CoV-2 E protein is expressed at a low level (20 *µ*M [*β*-D-1-thiogalactopyranoside]) in K^+^-uptake deficient bacteria. In this instance, inhibitory drugs reduce bacterial growth. The results in both panels may be compared to those obtained without any drug (black) or when the channel is uninduced (gray). The color scale indicates the different concentrations of the chemicals.

We recognize the potential of spurious factors to impact bacterial growth, leading to false identification of hits. Therefore, each chemical that scored positively in the negative assay was tested in a reciprocal assay, and in doing so, fallacious results are minimized significantly.

#### Positive assay

The second bacterial assay that we used is one in which the E protein is expressed at low levels in K^+^-uptake deficient bacteria. These bacteria are incapable of growth unless the media is supplemented by K^+^ (27). However, when a channel capable of K^+^ transport is heterologously expressed, the bacteria can thrive even in low K^+^ media (25, 26). Hence, in this instance, the viral channel **positively** impacts the bacteria, and channel blockers result in growth retardation. This scenario is entirely reciprocal to the negative assay described above.

Each of the hits identified in the negative assay was subjected to a dose-response analysis using the positive assay, as depicted in Figure 1b. The results present a mirror image of the negative assay, whereby in this instance the compounds decreased growth as follows: 5-Azacytidine (−29%), Plerixafor (−54%), Mebrofenin (−20%), Mavorixafor (trihydrochloride) (−20%), Plerixafor (octahydrochloride) (−49%), Cyclen (−27%), Kasugamycin (hydrochloride hydrate) (−72%), and Saroglitazar Magnesium (−26%). The values in parenthesis are the growth reduction of each chemical at 50 *µ*M relative to untreated bacteria. Detailed growth curves of each of the compounds can be found in Supplementary Figures 1-8.

#### Fluorescence-based test

The final test to examine the activity of channel blockers is based on detecting protein-mediated H^+^ flux. Bacteria that express a chromosomally-encoded pH-sensitive green fluorescent protein (28) exhibit fluorescence changes when their internal pH is altered. In particular, adding an acidic solution to the media will result in a readily detectable fluorescence change due to cytoplasmic acidification if the bacteria express a channel capable of H^+^ transport (29). Therefore, in this assay, blockers may be identified by their ability to diminish the fluorescence change.

As seen in Figure 2, most of the compounds are able to reduce the viroporin-induced fluorescence change with the exception of 5-Azacytidine and Mebrofenin. Saroglitazar was able to suppress the change entirely, while the impact of Cyclen was minor. Results at several drug concentrations including error bars can be found in Supplementary Figures 1-8.

**Fig. 2.**
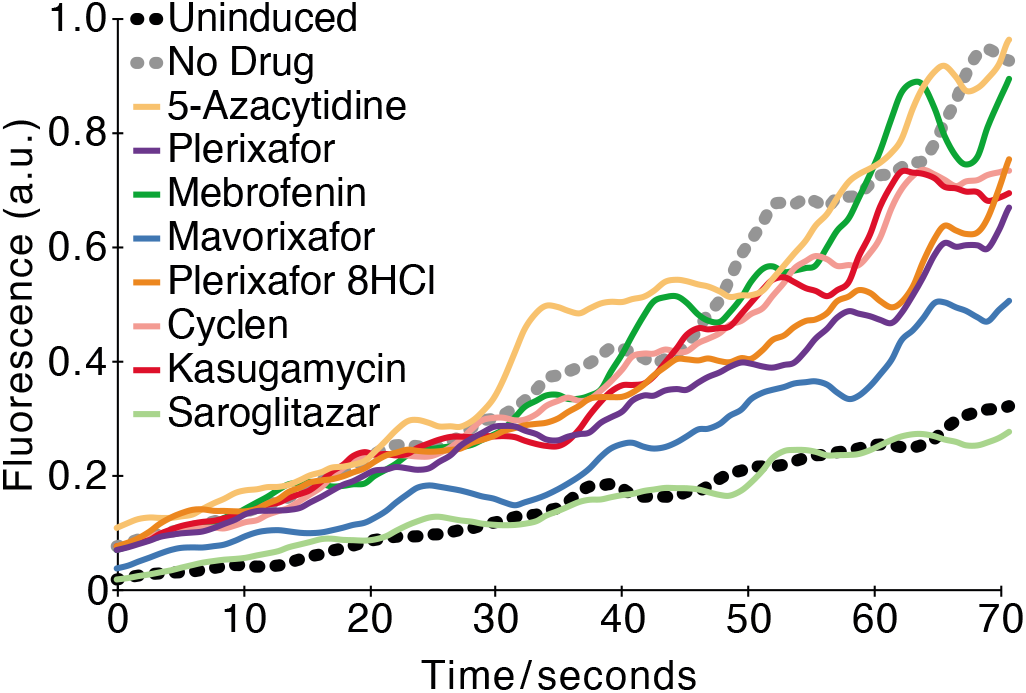
Fluorescence-based conductivity assay. The fluorescence of bacteria that harbor a pH sensitive GFP (28) and express the SARS-CoV-2 E protein was examined as a function of different chemicals at a concentration of 50 *µ*M. The experiment was performed as previously described (29), whereby at time 0, a concentrated solution of citric acid was injected into the media. The results may be compared to those obtained without any drug (black) or when the channel expression was not induced (gray).

### Antiviral activity

After completing the library screening, we tested the antiviral activity of each of the eight chemicals in Vero E6 tissue culture cells. Two additional compounds, gliclazide and memantine identified in a previous small screen (16) were also included.

The results shown in Figure 3a indicate that most of the chemicals are able to reduce viral replication as follows: 5-Azacytidine (−98%), Plerixafor (−53%), Mebrofenin (−45%), Mavorixafor (trihydrochloride) (−41%), Plerixafor (octahydrochloride) (−40%), Gliclazide (−41%), Cyclen (−38%), Kasugamycin (hydrochloride hydrate) (−27%), Saroglitazar Magnesium (−18%), and Memantine (−3%). The values in parenthesis are the growth reduction of each chemical at 10 *µ*M relative to untreated cells.

**Fig. 3.**
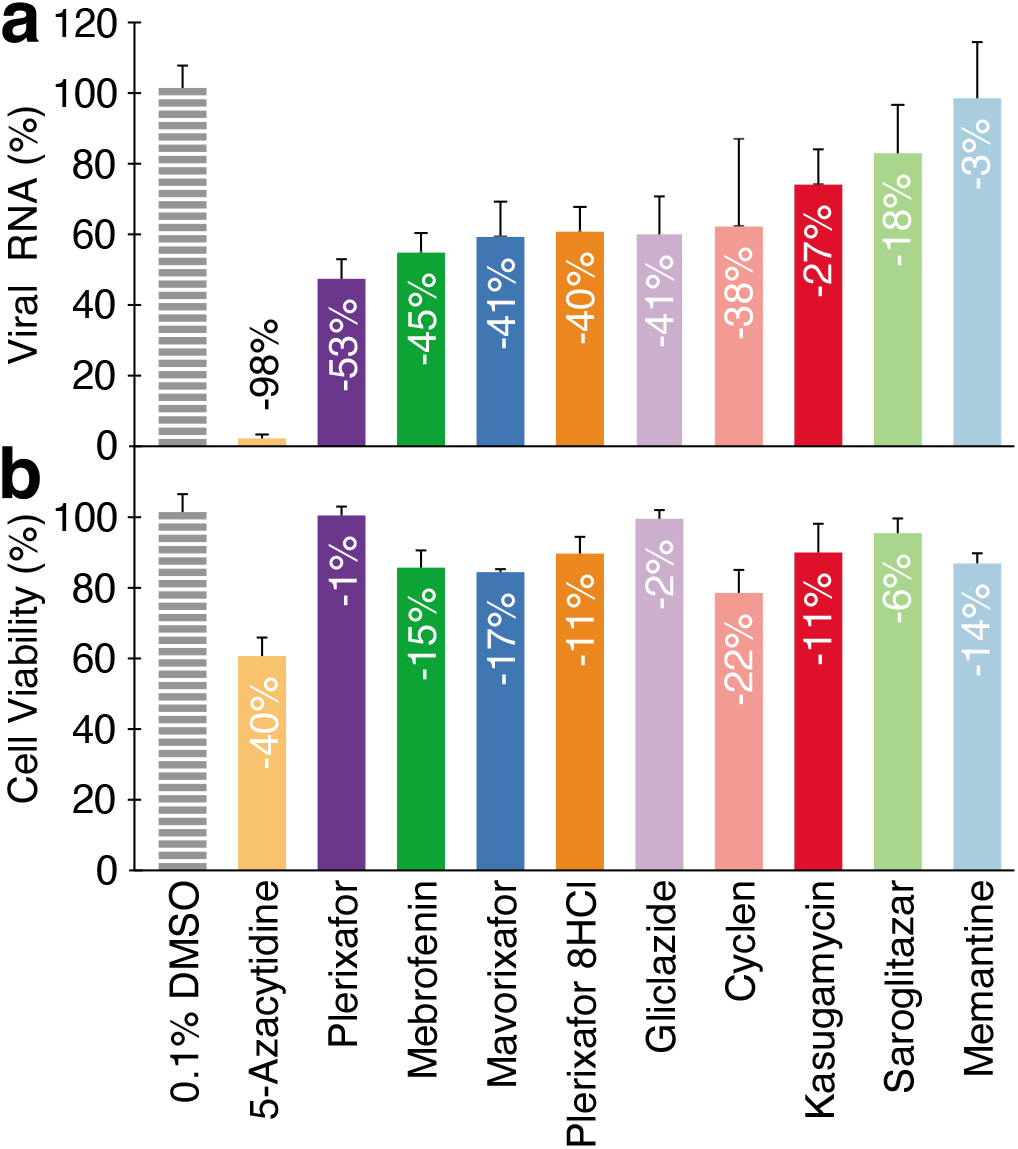
Tissue culture analysis of antiviral activity. a. The antiviral activity of each of the compounds at 10 *µ*M concentration was tested by its ability to minimize viral growth in Vero E6 cells as estimated by production of viral RNA. b. The toxicity of each of the compounds was measured using the 3-(4,5-dimethylthiazol-2-yl)-2,5-diphenyltetrazolium (MTT) assay (30). In both panels the results are compared to a blank treatment of 0.1% DMSO.

Finally, we estimated the the toxicity of each compound by assessing the cellular metabolic activity using a standard MTT assay (30), as shown in Figure 3b.

## Discussion

Repurposing has proven to be a reliable route towards drug discovery, in general, and for identifying antiviral drugs in particular. For example, the repurposing of azidothymidine (AZT) to combat AIDS (31, 32) was reported more than twenty years after its first description in 1964 (33).

Repurposing also represents one of the fastest approaches to curb infectivity (34). As an example, the only antiviral drug that is currently approved against COVID-19 is a product of repurposing - remdesivir. While its efficacy may still be a matter of contention (35–37), it is nonetheless an example of the speed at which drug repurposing can react to a health crisis.

Considering the above, it is not surprising that there have been numerous repurposing studies against SARS-CoV-2. While the vast majority have employed *in silico* screening, others have taken an experimental route. For example, Riva and co-workers screened 12,000 clinical-stage or FDA-approved drugs for their ability to inhibit viral replication (38). The encouraging results of this monumental study were 21 molecules that exhibited a dose-response activity profile.

Herein, a different and complementary approach was taken, focussing the screen on a single target of the virus - the E protein. Our rationale stemmed from the fact that channels are attractive drug targets, and searching for inhibitors against them is both rapid and economically viable in an academic setting. Furthermore, genetic selections in bacteria may cast a wider net when targeting an individual protein due to the host’s higher toxicity tolerance. Such studies also open the door to mutational analyses that may provide insight into protein function and drug resistance mechanisms (23).

Three independent bacteria-based assays were used to search the repurposed drug library. The first two tests are reciprocal, whereby in the negative assay, the channel is detrimental to bacterial growth, while in the positive assay, it is beneficial. Consequently, blockers will yield the opposite outcomes in both assays: In the negative assay, they will enhance growth, while in the positive assay, they will retard it. The use of two assays minimizes any erroneous hits: The negative assay is susceptible to any pleiotropic growth enhancers’ activity leading to false positives. Similarly, the positive assay would score a hit for any toxic compound. Yet, it is difficult to imagine how a drug can enhance the growth of bacteria in the negative assay while at the same time retard them in the positive assay if its effect was not specific. As an example, Plerixafor enhances bacterial growth fourfold in the negative assay and inhibits growth by sixfold in the positive assay.

The outcome of both tests comprised the list of compounds to be tested for antiviral activity. However, a third, fluorescence-based assay was employed to provide potential validation to the hits. In this test, a pH-sensitive GFP can report on the change of the cytoplasm’s acidity (28). Subsequently, the activity of a channel that is capable of H^+^ transport can be detected by measuring the fluorescence change due to acidification of the external media. Consequently, channel blockers would diminish the fluorescence change leading to their identification.

Gratifyingly, most of the hits identified by the positive and negative assays were able to lower the fluorescence change, except for 5-Azacytidine and Mebrofenin (Figure 2). Since non-specific factors influencing pH may obfuscate detection, we decided not to eliminate the latter two chemicals from our hits list to be as inclusive as possible.

In analyzing the outcomes of the screening tests, it is essential to realize that the assays are bacteria-based and, as such, should not be compared quantitatively to one another. Therefore, for screening purposes, any chemical that passed the positive and negative assay was pursued further. In addition we added to the list of hits obtained in the current study, gliclazide and memantine, identified from a previous small screen of focussed on channel blockers (16).

The results of the antiviral study in tissue culture were encouraging. Out of the ten chemicals in question, six reduced viral loads by 38-51%, two by 18-27%, and one by 98%. Memantine reduced viral RNA by a mere 3%, reflective of its low affinity (16). The different compounds also did not exhibit appreciable toxicity, except for 5-Azacytidine, a fact that may explain its extraordinary antiviral activity.

None of the compounds that the current screen yielded were identified by the large repurposing study of Riva and coworkers (38). One obvious factor that may explain the different outcomes between the two studies is stringency. Riva and coworkers screened every chemical at 5 *µ*M, whereas the current research in bacteria employed 50 *µ*M. Screening at this higher concentration stemmed from our desire to cast a wide net, which is feasible in the more tolerant bacterial system. While molecules can emerge from the bacterial screen with lower affinities, they may still be beneficial, serving as a starting point for detailed chemical exploration. Moreover, low-affinity drugs that block the E-channel may interact synergistically with inhibitors of other targets in the virus.

We recognize that while the compounds were retrieved by a screen searching for blockers of the E channel, they may impact the virus by a different route. This is particularly plausible for 5-Azacytidine that may rewire extensive transcriptional networks due to its influence on DNA methylation (39, 40).

In conclusion, we present a list of approved compounds for human use that inhibit SARS-CoV-2 to varying extents. Moreover, our approach may be used to screen rapidly for blockers of other viral ion channels in an effort to curb the virulence of their pathogenic host.

## Materials and Methods

### Channel assays

All three bacteria-based assays were conducted as described previously (16, 26).

In brief, over-night bacterial cultures were diluted and grown until their O.D.600 reached 0.2. 50 *µ*l of culture were subsequently transferred into 96-well flat-bottomed plates containing 50 *µ*l of the different treatments. Protein induction was achieved by adding *β*-D-1-thiogalactopyranoside at 100 *µ*M or 20 *µ*M for the negative and positive assays, receptively. D-glucose was added to a concentration of 1%. The plates were incubated for 16 hours at 37°C in a multi-plate incubator (Tecan Group, Männedorf, Switzerland) at a constant high shaking rate. O.D.600 readings were recorded every 15 min on a Infinite 200 plate reader (Tecan Group). For every measurement duplicates, or triplicates were conducted.

The positive assay was conducted in a similar manner except that the K^+^-uptake deficient bacteria were grown overnight and diluted in LB media in which Na^+^ was replaced by K^+^. Thereafter, the growth medium was replaced to LB that was supplemented with 5mM KCl.

The fluorescence-based assay was conducted with bacteria that harbor a chromosomal copy of a pH-sensitive GFP (29, 41). Overnight cultures were diluted 1:500 in LB media and grown up to an O.D.600 of 0.6-0.8. E protein expression was then induced by adding of 50 *µ*M *β*-D-1-thiogalactopyranoside to the growth media. After one hour of induction, the O.D.600 of all cells were measured, and after pelleting at 3500g for 10min, the bacteria were resuspended in McIlvaine buffer (200mM Na2HPO4, 0.9%NaCl adjusted to pH 7.6 with 0.1 M citric acid, 0.9%Nacl) to optical density of 0.25 at 600 nm. 200 *µ*l of cell suspension were subsequently transferred with 30 *µ*l of McIlvaine buffer to 96 well plate. The plate includes a row with only assay buffer and cultures without induction. The fluorescence measurement were carried out in a Infinite F200 pro microplate reader (Tecan Group, Männedorf, Switzerland).

At time zero, 70 *µ*l of 300mM Citric acid with 0.9% NaCl were added to the bacteria. The fluorescence emission of each well after addition of acid was measured by alternate read out of the two wavelengths for 30 seconds. The ratio for the two differently excited emissions, 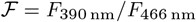 was calculated and translated into proton concentration according to (29, 41).

### Chemical screening

A library of 2839 repurposed drugs was purchased from MedChem Express (HY-L035, Monmouth Junction, NJ). Note that the number of chemicals in the library changes with time. Each chemical was added at 50 *µ*M concentration to the growth media with a total concentration of Dimethyl sulfoxide not exceeding 1%. All manipulations and growths were conducted on a Tecan EVO 75 robotic station (Männedorf, CH).

At first we screened all compounds in the negative assay using 96 well plates. Each plate had a positive and negative control. The positive control were bacteria in which channel expression was not induced, *i.e*., without *β*-D-1-thiogalactopyranoside. The negative control were bacteria to which DMSO was added without any chemicals.

Bacteria that exhibited growth enhancement above a certain empirical threshold were tested again in duplicate. Every compound that passed this test was then used in the positive assay in duplicate. Compounds that passed the positive and negative screen were then subjected to a dose-response analysis as well as a fluorescence-based study.

### Antiviral activity in tissue culture

#### Cells and viruses

Simian kidney Vero E6 (ATCC CRL-1586) cells were maintained in Eagle’s Minimum Essential Medium (EMEM; Biological Industries, Beit Haemek, Israel), supplemented with 10% fetal bovine serum, 2mM L-Glutamine, 10 IU/ml Penicillin, and 10 *µ*g/ml streptomycin (Biological Industries, Beit Haemek, Israel). SARS-CoV-2 clinical isolate (SARS-CoV-2 isolate Israel-Jerusalem-854/2020) was isolated on Vero E6 cells from a positive nasopharyngeal swab sample, obtained at the Hadassah Hospital Clinical Virology Laboratory. The virus was isolated and propagated (3 passages) in Vero E6 cells, and sequence verified. The virus titers of cleared infected cells supernatants were determined by a standard TCID50 assay on Vero E6 cells. Cell viability was monitored by the mitochondrial dehydrogenase enzyme (MTT) assay as previously described (42). All infection and tissue processing experiments were performed in a BSL-3 facility.

#### Antiviral susceptibility assay

Vero-E6 cells were pretreated for one hour with the tested compounds at 10 *µ*M, and were infected with SARS-CoV-2 at a multiplicity of infection (MOI) of 0.001-0.005 in the presence of the compounds (which were added to the culture at the time of viral inoculation, and further added during the culture incubation time). The medium with the same DMSO concentration was used as the no-drug control. Drug efficacies were assessed at 48 hours post infection by quantitation of the viral genomic RNA in the cell supernatants and the viral sub-genomic RNA within the infected cells.

#### RNA purification and quantification

Infected- and mock-infected cell cultures’ supernatants were flash-frozen and stored at −80°C until assayed. RNA was extracted using NucleoSpin RNA Mini kit for RNA purification (Macherey-Nagel, Cat #740955.250) according to the manufacturer’s instructions, and subjected to reverse transcription, using High-Capacity cDNA Reverse Transcription Kit (Thermo Fisher Scientific, Cat#). Quantitative real time (RT)-PCR was performed on a Quantstudio 3^™^ (Thermo Fisher Scientific) instrument, using TaqMan™ Fast Advanced Master Mix (Thermo Fisher Scientific, Cat# 4444558), with primers and probe targeted to the E gene.

## ACKNOWLEDGMENTS

This work was supported in part by grants from the Israeli Science Foundation, and the Israeli Science Ministry. The authors wish to thank Prof. Wolf and Ms. Hamdan from the Hadassah University Hospital for conducting the *in vitro* experiments, Prof. D. Engelberg and Dr. O. Cohen for technical assistance and useful discussions, and Prof. H Cedar for useful discussions. I.T.A. is the Arthur Lejwa Professor of Structural Biochemistry at the Hebrew University of Jerusalem.

