## supplemental figures for "Targeted Drug Repurposing Against the SARS-CoV-2 E channel Identifies Blockers With *in vitro* Antiviral Activity"

### Supplementary Information

**Prabhat Pratap Singh Tomar, Miriam Krugliak, and Isaiah Tuvia Arkin**[1]

Department of Biological Chemistry, The Alexander Silberman Institute of Life Sciences, The Hebrew University of Jerusalem, Edmond J. Safra Campus Givat-Ram, Jerusalem 91904, Israel

This manuscript was compiled on February 24, 2021

### Supplementary Figures

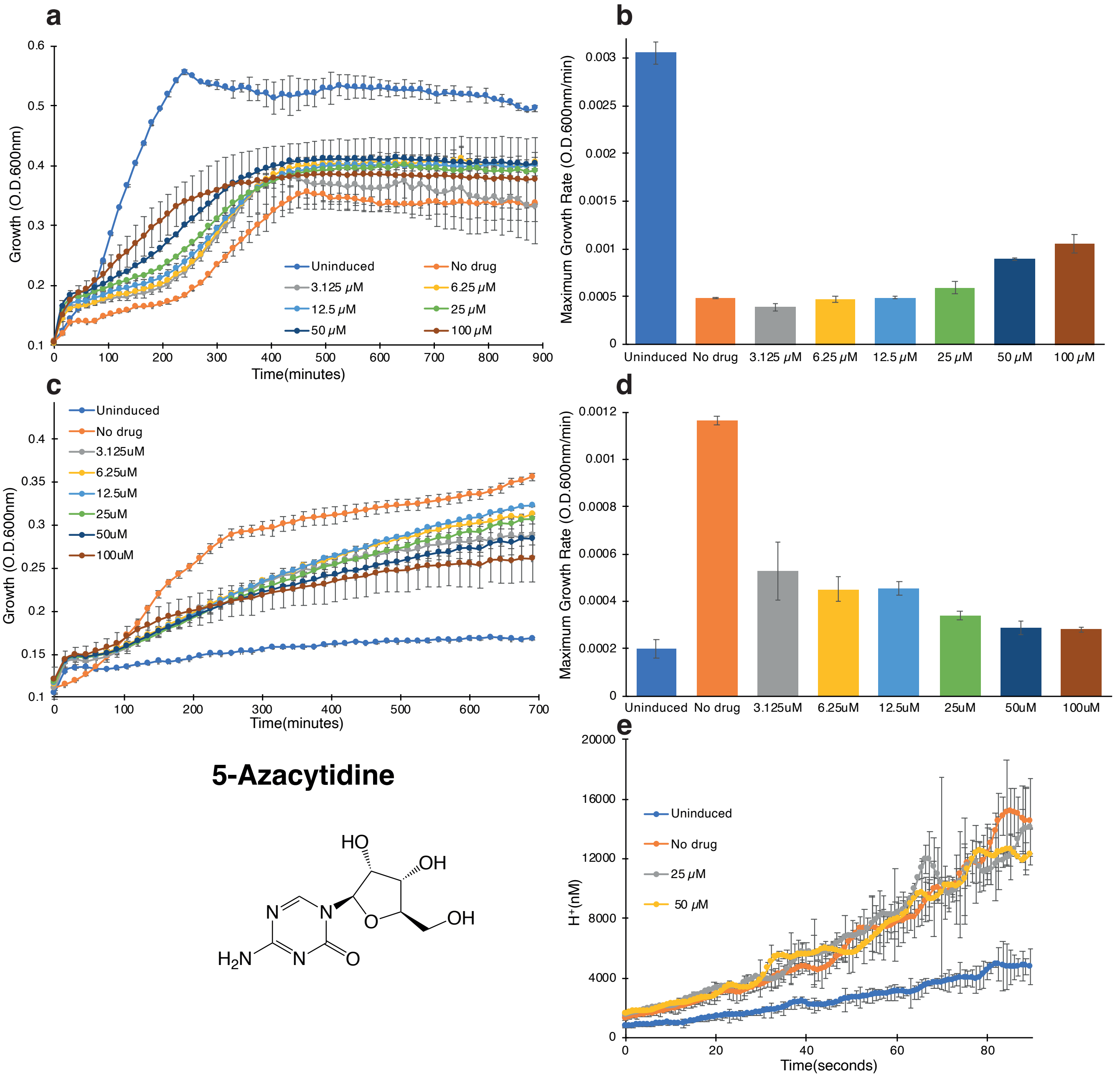


**Supplementary Fig. 1.** Raw screening data for 5-Azacytidine. a. Negative assay in which SARS-CoV-2 E protein is expressed at an elevated level (induced with 100 $\mu$M [$\beta$-D-1-thiogalactopyranoside]) and is therefore deleterious to bacteria. The different concentrations of the drug are indicated. b. Maximal growth rates obtained in the negative assay. c. Positive assay in which SARS-CoV-2 E protein is expressed at a low level (induced with 20 $\mu$M [$\beta$-D-1-thiogalactopyranoside]) in $K^+$-uptake deficient bacteria (1). In this instance, inhibitory drugs reduce bacterial growth. d. Maximal growth rates obtained in the positive assay. e. Fluorescence-based conductivity assay. The fluorescence of bacteria that harbor a pH sensitive GFP (2) and express the SARS-CoV-2 E protein was examined as a function of different chemical concnetration as noted. The experiment was performed as previously described (3), whereby at time 0, a concentrated solution of citric acid was injected into the media. In all panels LB indicates bacteria that do not express the channel as a positive control, while 100 $\mu$M IPTG indicates no drug added as a negative control.

The authors declare that thy are in the process of filing a patent for second medicinal use of Gliclazide and Memantine.

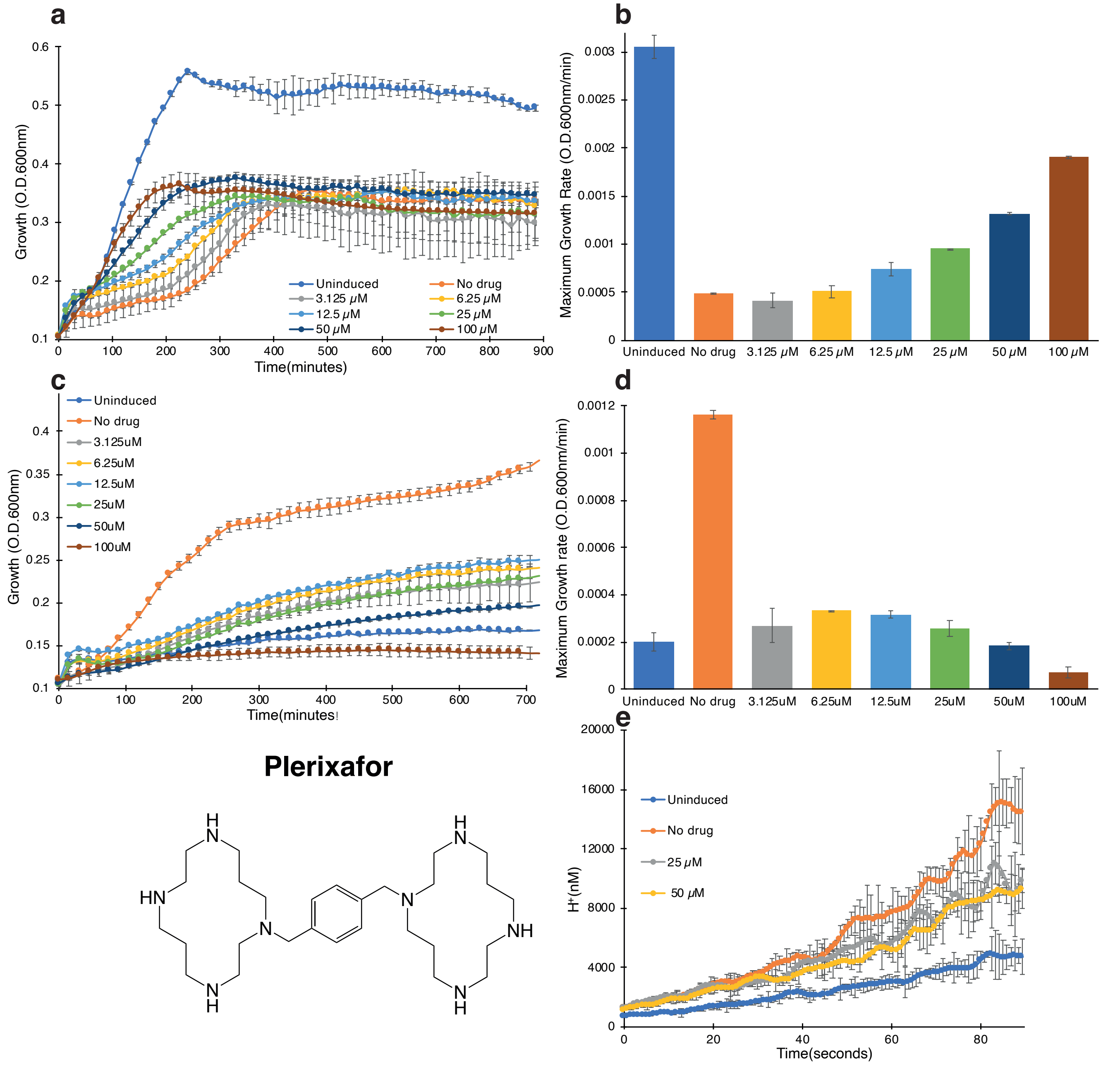


**Supplementary Fig. 2.** Raw screening data for Plerixafor. a. Negative assay in which SARS-CoV-2 E protein is expressed at an elevated level (induced with 100 $\mu$M [$\beta$-D-1-thiogalactopyranoside]) and is therefore deleterious to bacteria. The different concentrations of the drug are indicated. b. Maximal growth rates obtained in the negative assay. c. Positive assay in which SARS-CoV-2 E protein is expressed at a low level (induced with 20 $\mu$M [$\beta$-D-1-thiogalactopyranoside]) in $K^+$-uptake deficient bacteria (1). In this instance, inhibitory drugs reduce bacterial growth. d. Maximal growth rates obtained in the positive assay. e. Fluorescence-based conductivity assay. The fluorescence of bacteria that harbor a pH sensitive GFP (2) and express the SARS-CoV-2 E protein was examined as a function of different chemical concnetration as noted. The experiment was performed as previously described (3), whereby at time 0, a concentrated solution of citric acid was injected into the media. In all panels LB indicates bacteria that do not express the channel as a positive control, while 100 $\mu$M IPTG indicates no drug added as a negative control.

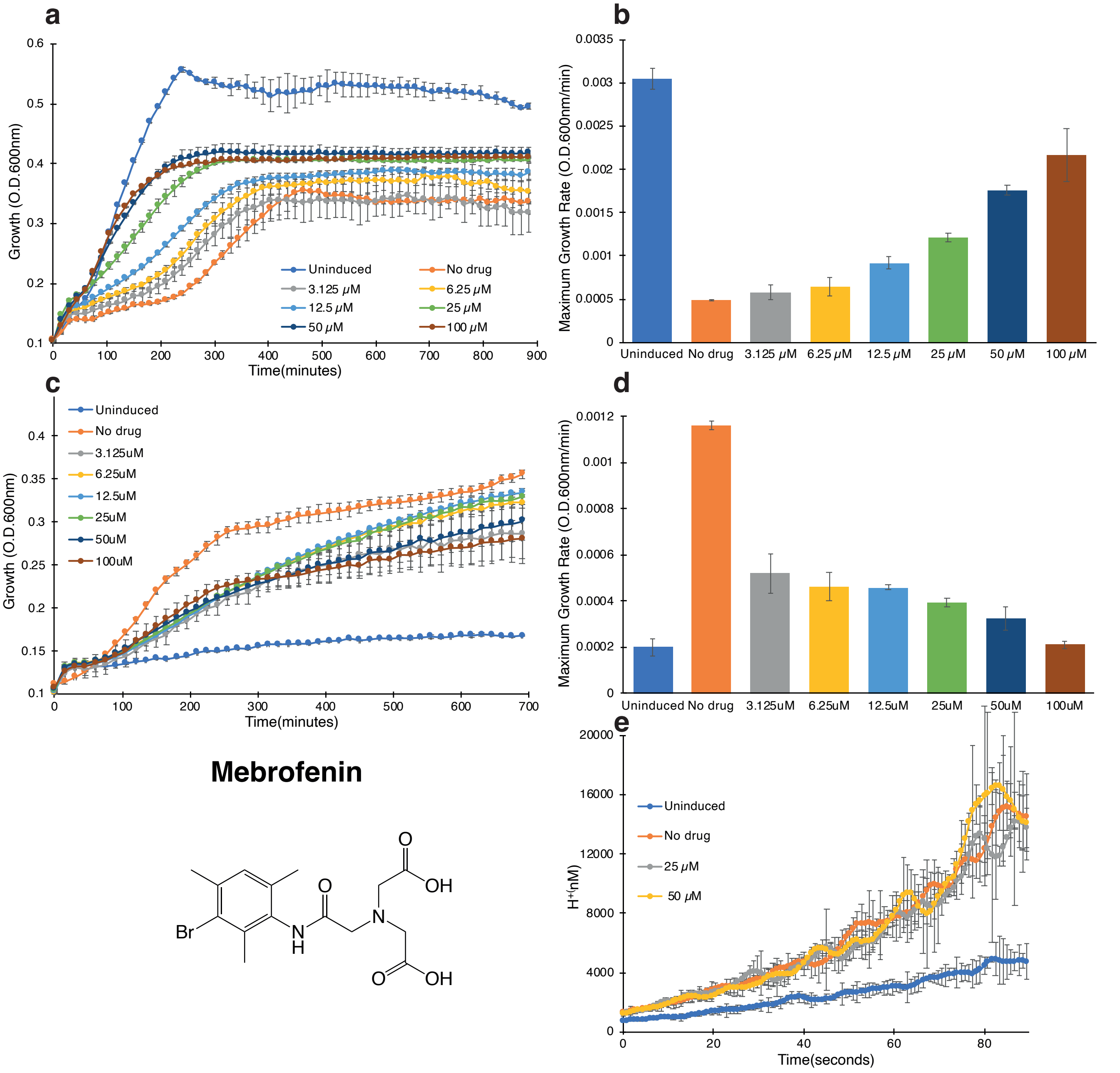


**Supplementary Fig. 3.** Raw screening data for Mebrofenin. a. Negative assay in which SARS-CoV-2 E protein is expressed at an elevated level (induced with 100 $\mu$M [$\beta$-D-1-thiogalactopyranoside]) and is therefore deleterious to bacteria. The different concentrations of the drug are indicated. b. Maximal growth rates obtained in the negative assay. c. Positive assay in which SARS-CoV-2 E protein is expressed at a low level (induced with 20 $\mu$M [$\beta$-D-1-thiogalactopyranoside]) in $K^+$-uptake deficient bacteria (1). In this instance, inhibitory drugs reduce bacterial growth. d. Maximal growth rates obtained in the positive assay. e. Fluorescence-based conductivity assay. The fluorescence of bacteria that harbor a pH sensitive GFP (2) and express the SARS-CoV-2 E protein was examined as a function of different chemical concnetration as noted. The experiment was performed as previously described (3), whereby at time 0, a concentrated solution of citric acid was injected into the media. In all panels LB indicates bacteria that do not express the channel as a positive control, while 100 $\mu$M IPTG indicates no drug added as a negative control.

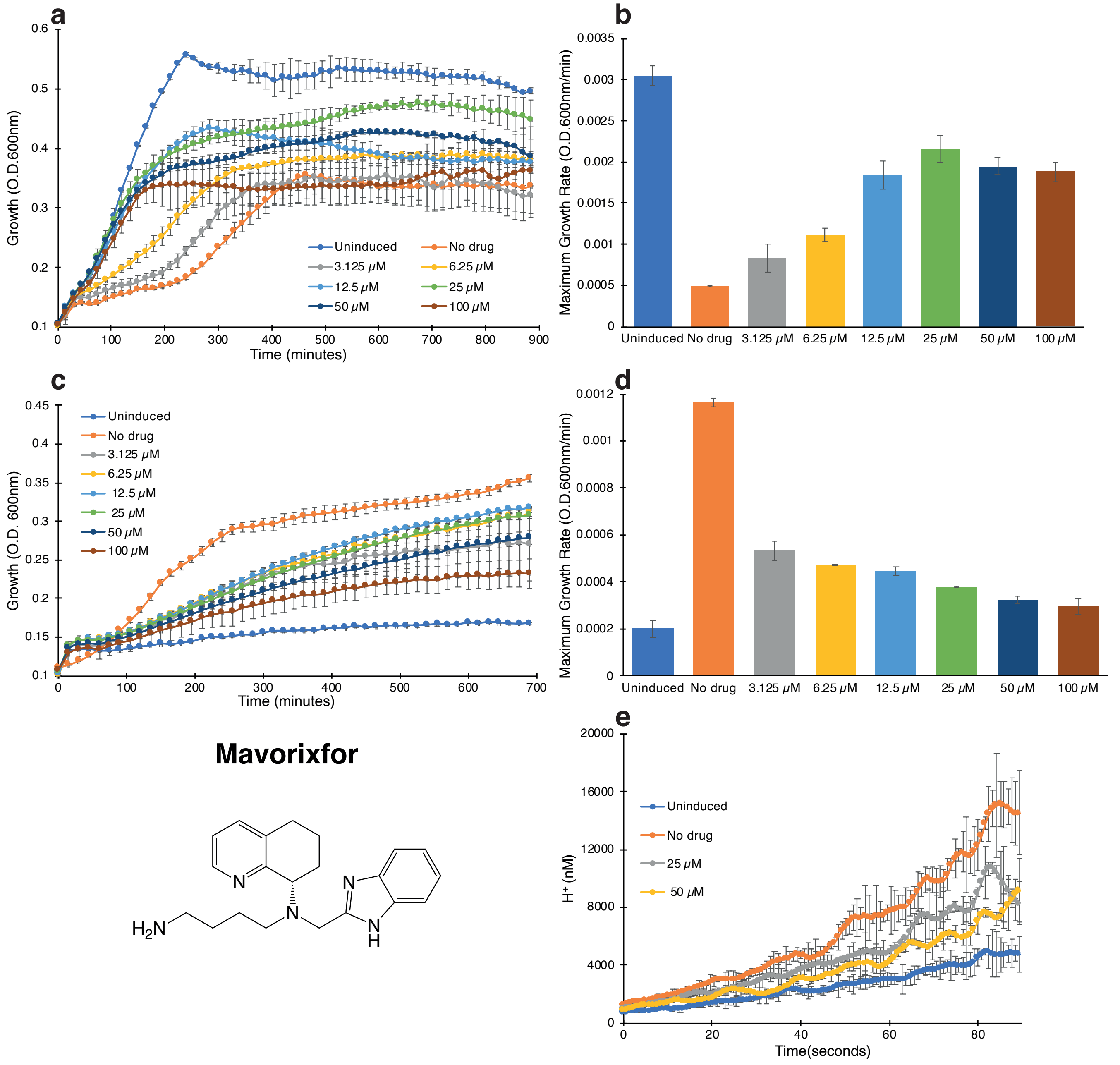


**Supplementary Fig. 4.** Raw screening data for Mavorixafor. a. Negative assay in which SARS-CoV-2 E protein is expressed at an elevated level (induced with 100 $\mu$M [$\beta$-D-1-thiogalactopyranoside]) and is therefore deleterious to bacteria. The different concentrations of the drug are indicated. b. Maximal growth rates obtained in the negative assay. c. Positive assay in which SARS-CoV-2 E protein is expressed at a low level (induced with 20 $\mu$M [$\beta$-D-1-thiogalactopyranoside]) in $K^+$-uptake deficient bacteria (1). In this instance, inhibitory drugs reduce bacterial growth. d. Maximal growth rates obtained in the positive assay. e. Fluorescence-based conductivity assay. The fluorescence of bacteria that harbor a pH sensitive GFP (2) and express the SARS-CoV-2 E protein was examined as a function of different chemical concnetration as noted. The experiment was performed as previously described (3), whereby at time 0, a concentrated solution of citric acid was injected into the media. In all panels LB indicates bacteria that do not express the channel as a positive control, while 100 $\mu$M IPTG indicates no drug added as a negative control.

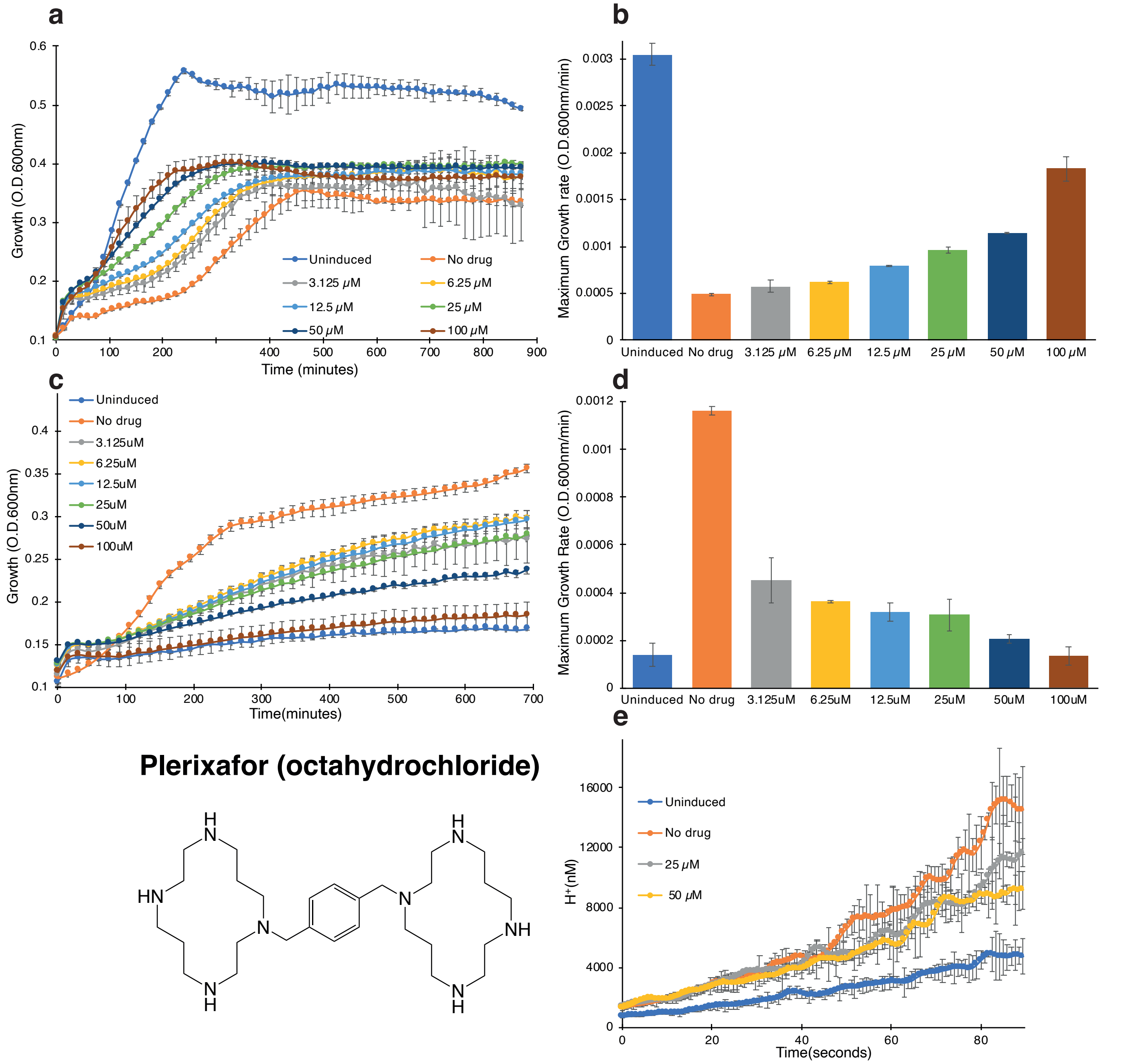


**Supplementary Fig. 5.** Raw screening data for Plerixafor (octahydrochloride). a. Negative assay in which SARS-CoV-2 E protein is expressed at an elevated level (induced with 100 $\mu$M [$\beta$-D-1-thiogalactopyranoside]) and is therefore deleterious to bacteria. The different concentrations of the drug are indicated. b. Maximal growth rates obtained in the negative assay. c. Positive assay in which SARS-CoV-2 E protein is expressed at a low level (induced with 20 $\mu$M [$\beta$-D-1-thiogalactopyranoside]) in $K^+$-uptake deficient bacteria (1). In this instance, inhibitory drugs reduce bacterial growth. d. Maximal growth rates obtained in the positive assay. e. Fluorescence-based conductivity assay. The fluorescence of bacteria that harbor a pH sensitive GFP (2) and express the SARS-CoV-2 E protein was examined as a function of different chemical concnetration as noted. The experiment was performed as previously described (3), whereby at time 0, a concentrated solution of citric acid was injected into the media. In all panels LB indicates bacteria that do not express the channel as a positive control, while 100 $\mu$M IPTG indicates no drug added as a negative control.

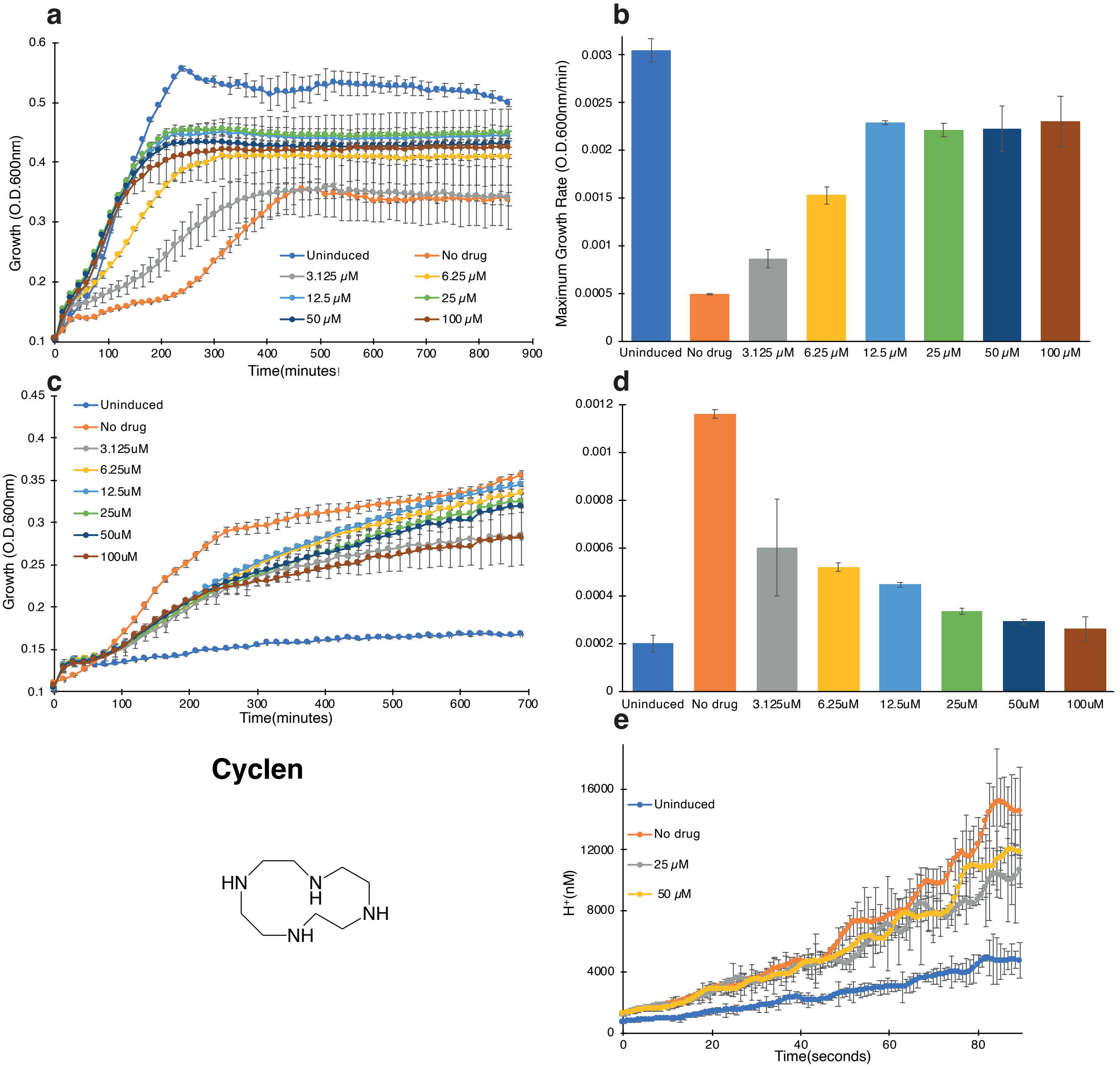


**Supplementary Fig. 6.** Raw screening data for Cyclen. a. Negative assay in which SARS-CoV-2 E protein is expressed at an elevated level (induced with 100 $\mu$M [$\beta$-D-1-thiogalactopyranoside]) and is therefore deleterious to bacteria. The different concentrations of the drug are indicated. b. Maximal growth rates obtained in the negative assay. c. Positive assay in which SARS-CoV-2 E protein is expressed at a low level (induced with 20 $\mu$M [$\beta$-D-1-thiogalactopyranoside]) in $K^+$-uptake deficient bacteria (1). In this instance, inhibitory drugs reduce bacterial growth. d. Maximal growth rates obtained in the positive assay. e. Fluorescence-based conductivity assay. The fluorescence of bacteria that harbor a pH sensitive GFP (2) and express the SARS-CoV-2 E protein was examined as a function of different chemical concnetration as noted. The experiment was performed as previously described (3), whereby at time 0, a concentrated solution of citric acid was injected into the media. In all panels LB indicates bacteria that do not express the channel as a positive control, while 100 $\mu$M IPTG indicates no drug added as a negative control.

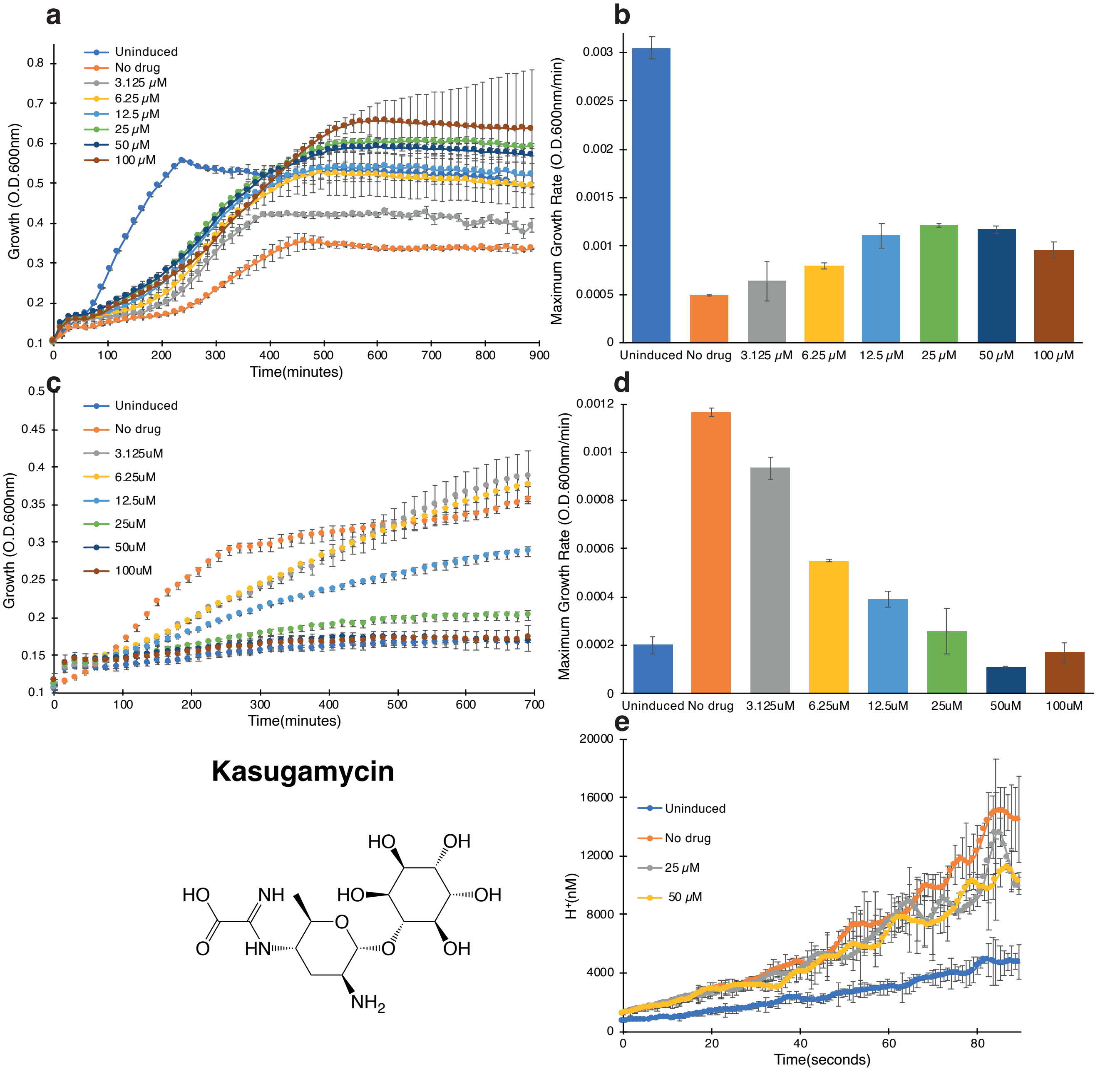


**Supplementary Fig. 7.** Raw screening data for Kasugamycin. a. Negative assay in which SARS-CoV-2 E protein is expressed at an elevated level (induced with 100 $\mu$M [$\beta$-D-1-thiogalactopyranoside]) and is therefore deleterious to bacteria. The different concentrations of the drug are indicated. b. Maximal growth rates obtained in the negative assay. c. Positive assay in which SARS-CoV-2 E protein is expressed at a low level (induced with 20 $\mu$M [$\beta$-D-1-thiogalactopyranoside]) in $K^+$-uptake deficient bacteria (1). In this instance, inhibitory drugs reduce bacterial growth. d. Maximal growth rates obtained in the positive assay. e. Fluorescence-based conductivity assay. The fluorescence of bacteria that harbor a pH sensitive GFP (2) and express the SARS-CoV-2 E protein was examined as a function of different chemical concnetration as noted. The experiment was performed as previously described (3), whereby at time 0, a concentrated solution of citric acid was injected into the media. In all panels LB indicates bacteria that do not express the channel as a positive control, while 100 $\mu$M IPTG indicates no drug added as a negative control.

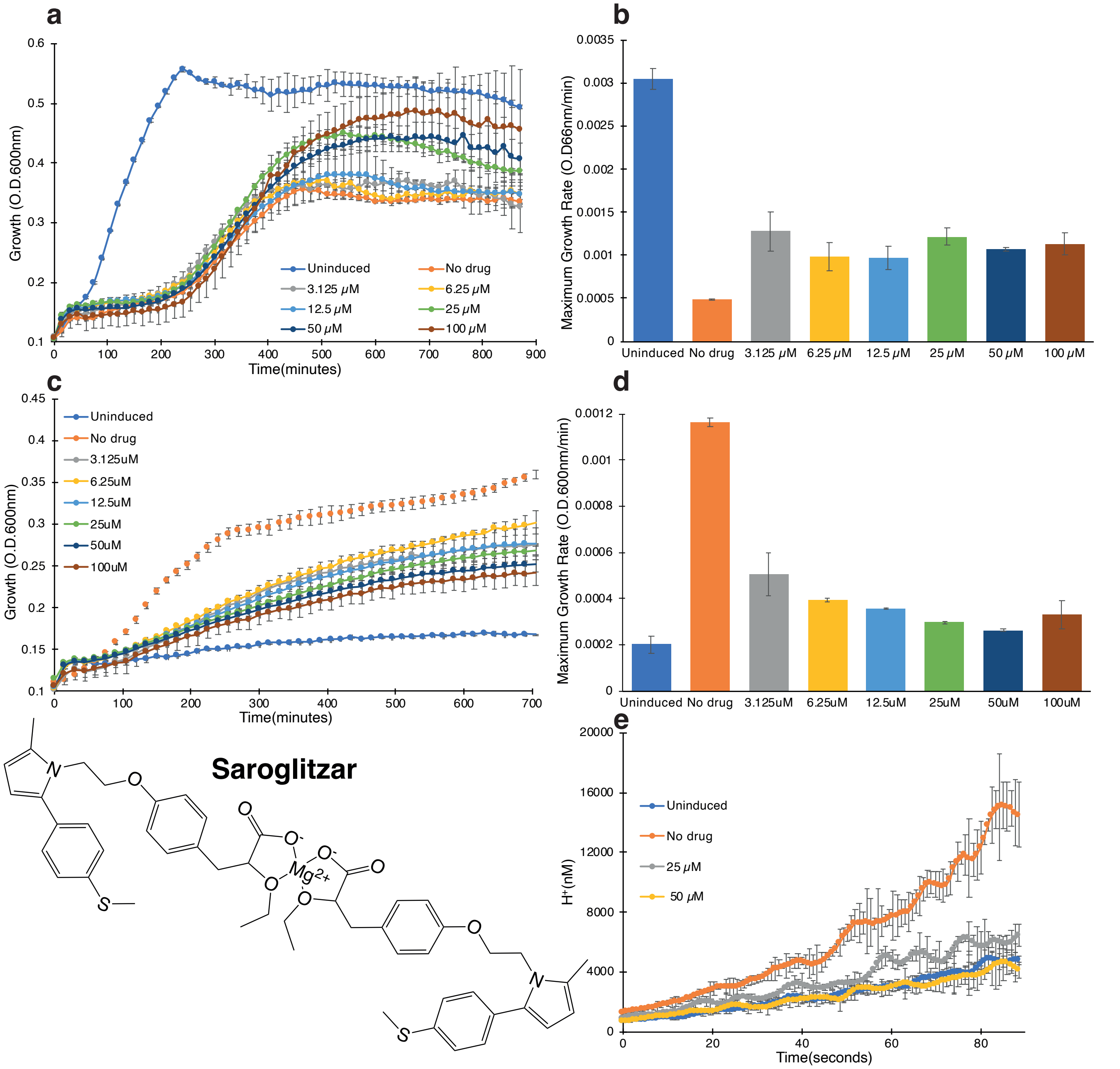


**Supplementary Fig. 8.** Raw screening data for Saroglitazar. a. Negative assay in which SARS-CoV-2 E protein is expressed at an elevated level (induced with 100 $\mu$M [$\beta$-D-1-thiogalactopyranoside]) and is therefore deleterious to bacteria. The different concentrations of the drug are indicated. b. Maximal growth rates obtained in the negative assay. c. Positive assay in which SARS-CoV-2 E protein is expressed at a low level (induced with 20 $\mu$M [$\beta$-D-1-thiogalactopyranoside]) in $K^+$-uptake deficient bacteria (1). In this instance, inhibitory drugs reduce bacterial growth. d. Maximal growth rates obtained in the positive assay. e. Fluorescence-based conductivity assay. The fluorescence of bacteria that harbor a pH sensitive GFP (2) and express the SARS-CoV-2 E protein was examined as a function of different chemical concnetration as noted. The experiment was performed as previously described (3), whereby at time 0, a concentrated solution of citric acid was injected into the media. In all panels LB indicates bacteria that do not express the channel as a positive control, while 100 $\mu$M IPTG indicates no drug added as a negative control.
